# Differences and similarities between human hippocampal low-frequency oscillations during navigation and mental simulation

**DOI:** 10.1101/2024.12.04.626897

**Authors:** Sarah Seger, Jennifer Kriegel, Brad Lega, Arne Ekstrom

**Affiliations:** Neuroscience Interdisciplinary Program, University of Arizona, 1503 E. University Blvd., Tucson, AZ 85719; Department of Neurosurgery, University of Texas Southwestern Medical School, Dallas, TX; Psychology Department, University of Arizona, 1503 E. University Blvd., Tucson, AZ 85719; Evelyn McKnight Brain Institute, University of Arizona, 1503 E. University Blvd., Tucson, AZ 85719

## Abstract

Low frequency oscillations in the hippocampus emerge during by both spatial navigation and episodic memory function in humans. We have recently shown that in humans, memory-related processing is a stronger driver of low frequency oscillations than navigation. These findings and others support the idea that low-frequency oscillations are more strongly associated with a general memory function than with a specific role in spatial navigation. However, whether the low-frequency oscillations that support episodic memory and those during navigation could still share some similar functional roles remains unclear. In this study, patients undergoing intracranial electroencephalography (iEEG) monitoring performed a navigation task in which they navigated and performed internally directed route replay, similar to episodic memory. We trained a random forest classification model to use patterns in low-frequency power (2-12 Hz) to learn the position during navigation and subsequently used the same model to successfully decode position during mental simulation. We show that removal of background differences in power between navigation and mental simulation is critical to detecting the overlapping patterns. These results suggest that the low-frequency oscillations that emerge during navigation are more associated with a role in memory than specifically with a navigation related function.

## Introduction

Studies suggest that the hippocampus contributes to both episodic memory and spatial navigation but the mechanistic basis for these contributions remains unclear. An important contributor to navigation and episodic memories is the emergence of low-frequency (2 – 12 Hz) oscillations (LFOs) in the local field potential of the hippocampus, also called “theta” oscillations. Theta oscillations have been most thoroughly studied in the rodent hippocampus where their emergence is strongly linked to voluntary, goal-directed movement (McFarland et al., 1975; Czurkó et al., 1999; Sheremet et al., 2016; Kropff et al., 2021; Kennedy et al., 2022) One long-standing model based on this observation argues that theta oscillations are related to the goal-directed movement and sensorimotor integration that occurs during navigation (Bland and Oddie, 2001; Lakatos et al., 2008; Schroeder et al., 2010) Studies with human patients undergoing intracranial monitoring observe increases in theta oscillations during movement when navigating in a virtual environment on desktop computer, which also supports a role for theta oscillations and sensorimotor activity and coding of movement-related variables (Ekstrom et al., 2005, 2009; Watrous et al., 2011; Aghajan et al., 2017; Bohbot et al., 2017; Bush et al., 2017) Notably, the theta oscillations that emerge during movement in humans are more transient, occurring in brief bouts as opposed to being sustained, as compared to those observed in rodents (Watrous et al., 2013).

Robust theta oscillations are also observed during the encoding and retrieval periods of verbal memory tasks. Memory encoding and retrieval has no overt demands on movement, sensorimotor integration, or spatial component (Sederberg et al., 2003; Lega et al., 2012; Solomon et al., 2017). Given that memory is engaged during navigation, one possibility then is that the theta oscillations that emerge during navigation serve a memory-related function rather than a direct sensorimotor related function (Eichenbaum, 2017a). If the theta oscillations that emerge during navigation are memory related, then this predicts that the theta oscillations which emerge in subsequent navigation-related memory retrieval should show some similarity to those present during navigation.

We previously showed that the hippocampal low-frequency oscillations during mental simulation are significantly higher in power, frequency, and duration compared to navigation (Seger et al., 2023). While this suggests differences in the theta oscillations during navigation and internally generated memory processing, it is unclear whether there is overlap in the navigation related informational content or functional role of these theta oscillations. Previous papers have reported evidence for a compressed reactivation of patterns present at encoding during retrieval (Michelmann et al., 2019; Wimmer et al., 2020). These studies, however, employed either scalp EEG or MEG and therefore could not definitively relate the measured signals to hippocampal low-frequency oscillations. Additionally, iEEG studies in humans have found that low-frequency oscillatory power present in the MTL during encoding of word pairs are reinstated at upon retrieval following the presentation of a partial cue (Yaffe et al., 2014).The presence of a partial cue at memory retrieval, however, makes it difficult to separate the contribute of external sensory input from memory related activity. Moreover, because studies of episodic memory often involve one-off learning, the nature of underlying information represented by these oscillations is difficult to determine To test whether there is evidence for a similar functional role of theta oscillations during navigation and internally generated memory processing, it is necessary to use a paradigm in which route replay is internally driven (i.e. entirely memory driven) but can be behaviorally validated and compared with navigation. To this end, we designed a virtual navigation study in which participants used a joystick to navigate between target stores and subsequently mentally simulated the traversed routes in the absence of any external cues. We recorded electrical activity from the hippocampus as human patients undergoing invasive monitoring for epilepsy treatment performed the experimental task. We first test whether position-related information could be identified in the patterns of low-frequency power during navigation; we then used this information to determine whether position could also be decoded during subsequent route replay and if so, under what conditions.

## Results

iEEG patients performed a spatial navigation task on a laptop computer using an Xbox controller. As shown in Figure 1a, the spatial navigation task required participants to use a joystick to navigate between stores within a sparse environment (Figure 1a) and subsequently perform mental simulation by “mentally navigating” the same route. Mental simulation of each route began once the subject arrived at the storefront of the target destination and occurred as patients stared at a black screen with a white crosshair to limit eye movement (i.e., in the absence of sensory input). This occurred on most trials without movement of the joystick except on catch trials, which ensured that patients were simulating the routes throughout (Figure 1a; see Methods).

**Figure 1:**
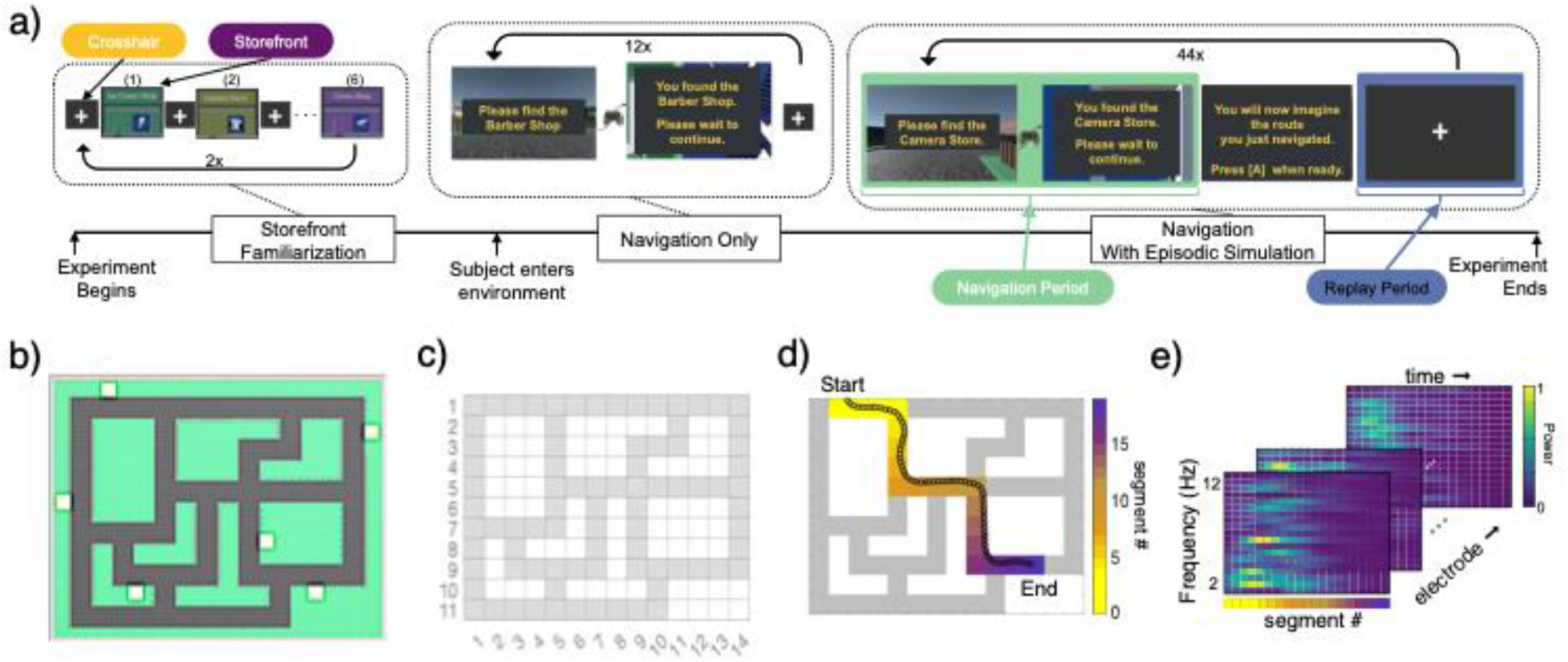
Navigation task design, behavioral results, and segmentation for decoding training data. a. Schematic of the experimental design. Store Familiarization– static images of each storefront were presented for 5000ms on a black background and followed by 5000ms of crosshair presentation; each storefront was presented 2x. Navigation Only– the subject was placed outside 1 of the 6 storefronts and used the XBox controller to locate the target store that was shown on the screen prompt (target store names remained visible at all times). Upon arrival at the proper target store, a 5000ms crosshair was presented before a new prompt appeared for the next target. The participants navigated to each store 2x. Navigation + Mental Simulation– similar to navigation only except, upon arrival at the proper target store, an on-screen prompt instructed participants to mentally navigate by picturing themselves at the starting location. Once prepared, the participants pressed [A] using the controller to start mental simulation and again once they imagined themselves outside the target destination. b. Environment layout showing road configuration and location of the 6 target stores. c. Segmentation of environment into 82 spatial bins with size 10 vm^2^. d. Example trajectory for a single subject for a single navigation trial. The participant began navigation at the store located on the top left of the environment (yellow colors) and navigated to the store at the bottom right of the environment (purple colors). Each time point during mobile periods of navigation was assigned to spatial bins based on the position. e. Example 2- 12 Hz power for all hippocampal electrodes for the subject for each spatial bin traversed for the trajectory shown in d. Each ‘sheet’ represents a unique electrode.

### Raw low-frequency power from hippocampal electrodes can be used to decode position during navigation

To determine whether there is sufficient information embedded in low-frequency (2 – 12 Hz) oscillatory power to decode position during navigation, the spatial environment (Figure 1b) was divided into 82 equally sized segments (Figure 1c). For every navigation trial, we assigned each time point to a spatial segment based on the position and subsequently averaged across time points assigned to each individual segment. For each patient, we combined the average low frequency power for each segment across all trials and all electrodes to generate a single matrix per patient which contained the low-frequency power for all electrodes and every segment that was traversed throughout all navigation trials (see Methods).

We trained a random forest classification ensemble (see Methods) to learn position using the average low frequency power (2- 12 Hz) from all hippocampal electrodes plus the identity of the start and end store as predictor variables. To compute chance accuracy, we trained an identical version of the model on training data with segment labels that were shuffled. Across participants, the mean accuracy of the model trained for decoding position during navigation using the true segment labels was significantly greater than mean accuracy of the model trained with shuffled segment labels (Wilcoxon ranksum test p = 0.014, median true labels = 0.033, median shuffle labels = 0.022, Figure 2a). The mean accuracy for each of the 82 spatial segments is shown in Figure 2b. According to an analysis of feature of importance, the frequency with the greatest predictive value was about 4.8 Hz (Figure 2c).

**Figure 2:**
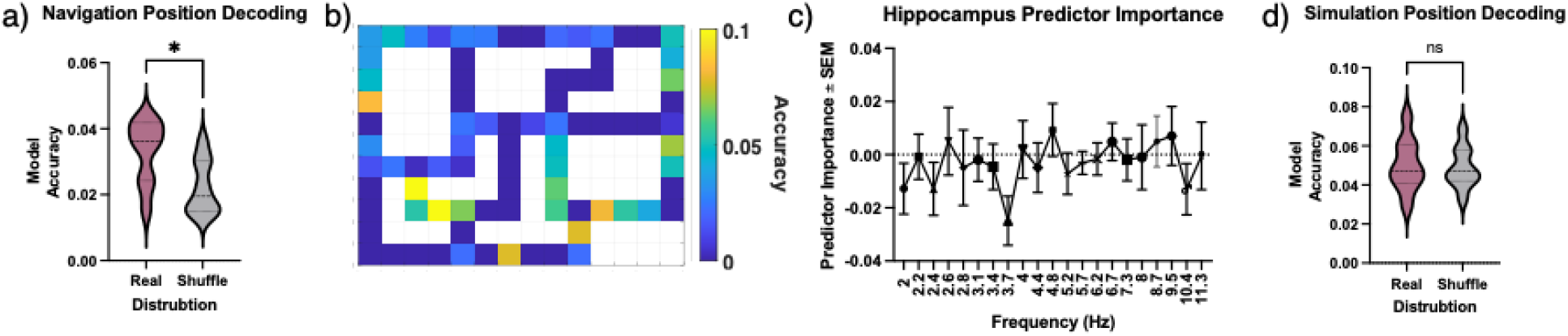
Position decoding during is successful during navigation and unsuccessful during mental simulation when using the raw power values. a. Across subjects, the mean accuracy of position decoding during navigation for the true segment labels was significantly greater than mean accuracy of the model trained with shuffled segment labels. b. Heat map showing the mean accuracy for each spatial segment. c. Average feature importance for all frequencies between 2- 12 Hz. d. Across subjects, the mean accuracy of position decoding during mental simulation using the model trained during navigation was not significantly greater than mean accuracy of the model trained on the shuffled time series.

To determine whether the patterns in power for frequencies above 12 Hz contain position related information, we trained a version of the model using the power for log-spaced frequencies between 2 and 100Hz. When we included all frequencies between 2 – 100Hz, the ability to decode position during navigation was still significantly greater than expected by chance (Supplemental Figure 1a). There was no difference in the decoding accuracy, however, for the model trained on 2- 12Hz compared to the model trained on 2- 100 Hz (Supplemental Figure 1d). Consistent with our feature importance ranking (Figure 2c), this suggests that positional information in the local field potential is primarily present within the low-frequency delta-theta-alpha band.

### Patterns of low-frequency power in the raw local field potential during navigation cannot be used to decode position during mental simulation

We next tested whether there is overlap in the patterns embedded in the low frequency local field potential between navigation and mental simulation. To test the hypotheses that there is overlap between the representation of position during navigation and mental simulation, we used the model trained on low-frequency power during navigation to decode the position at each time point during mental simulation. To do this, we first down-sampled the position during each navigation trial to match the duration of mental simulation for the same route. This was necessary because mental simulation happened at a compressed rate compared to navigation: typically, about 2-3 times the rate of navigation. This yielded an expected position for each time point during mental simulation. For every time point during each mental simulation trial, we compared the expected position to the position that was predicted by the model. For each subject, we then computed the average number of correctly identified time points for each trial and averaged across all trials. To determine chance accuracy, we shuffled the time points during each mental simulation trial and applied the same model trained during navigation to the power at each time point of the shuffled time series.

For the model trained on raw power during navigation and applied to the raw power during mental simulation, the mean decoding accuracy did not significantly differ between the real and shuffled timeseries (Wilcoxon ranksum test p = 0.180, accuracy true timeseries = 0.050, accuracy shuffled timeseries = 0.048, Figure 2d). We also could not decode position during mental simulation using the model trained during navigation when we included all frequencies between 2 – 100Hz (Supplemental Figure 1c). Therefore, although we could decode position from navigation using the raw power recorded from navigation, this did not generalize to mental simulation. There are some likely important reasons for this null finding, however. As we noted in our previous paper, simulation and navigation differ in terms of power and frequency, with both being higher for simulation. Therefore, the differences in power and frequency could have overshadowed more subtle memory related patterns shared between navigating the same route and simulating it.

### Navigation and mental simulation are dissociable based on the raw local field potential

To better understand why we might not be able to decode the contents of simulation from the raw power recorded during navigation of the same route – despite them involving overlapping encoded and retrieved memories, we wished to determine whether the raw low frequency power during navigation and simulation was sufficient to decode the condition (navigation or mental simulation). This would suggest that there might be important differences between the two conditions that a classifier could pick up on. To do this, we first found the average low frequency power over a randomly selected 1 second interval for each navigation and mental simulation trial for all electrodes. For each patient, we trained a random forest classification ensemble to use the low frequency power from all electrodes to differentiate between navigation and mental simulation conditions. To compute chance accuracy, we randomly shuffled the condition labels and trained an identical version of the model on the shuffled dataset.

When using the raw power during navigation and mental simulation, we were able to decode navigation from mental simulation at a level that was significantly greater than expected by chance (Wilcoxon ranksum test p = 0.027, mean true accuracy = 0.540, mean shuffle accuracy = 0.470, Figure 3a). As shown in the supplemental figures, we were also able to decode other differences between navigation and simulation, such as start periods, stop periods, and other epochs that otherwise might have shared similarities between navigation and simulation (Supplemental Figure 2 a -h).This suggests that the difference in power (and frequency) between navigation and mental simulation might have been obscuring subtle patterns in the local field potential that might reveal embedded memory signals in the local field potential.

**Figure 3:**
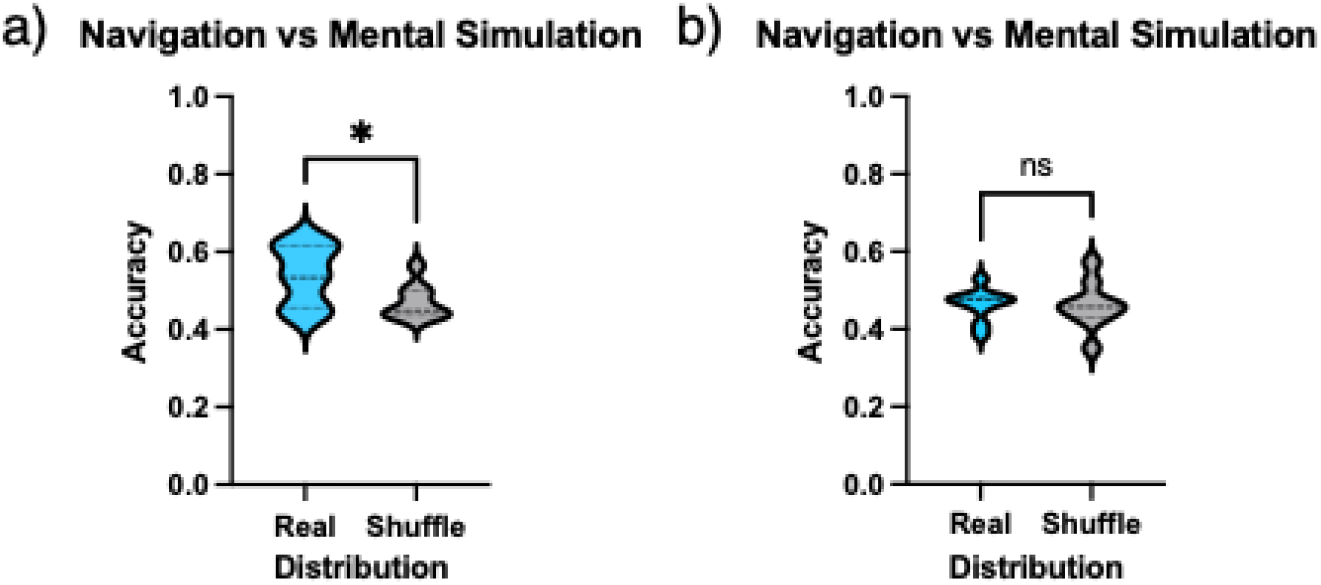
Raw but not standardized power can be used to decode the navigation from the mental simulation condition a. When using the raw low-frequency power, the mean accuracy for decoding navigation versus mental simulation condition for the model trained with true condition labels was significantly greater than the model trained with shuffled condition labels across all subjects. b. When using the standardized low-frequency power, the mean accuracy for decoding navigation versus mental simulation condition for the model trained with true condition labels was not significantly different than the model trained with shuffled condition labels across all subjects.

Given that we could readily decode epochs of navigation from those of mental simulation, we reasoned that better equating them might reveal any shared memory signals between the two. One way to reduce the differences in the underlying power spectrum between navigation and mental simulation is by standardizing the power with each condition. We standardized the power within each condition and each frequency on an electrode-by-electrode basis by subtracting the mean and dividing by the standard deviation (see Methods). To test whether standardizing the power within each condition removes differences in the underlying power spectrum between navigation and mental simulation, we trained a new random forest classification ensemble to differentiate standardized power in a manner identical to how we had applied the classifier for raw power. When using the standardized power during navigation and mental simulation, we were not able to decode navigation from mental simulation at a level significantly greater than expected by chance (Wilcoxon ranksum test p = 0.500, mean accuracy =0.456, mean shuffle accuracy = 0.453, Figure 3b). This suggested that the differences that had earlier defined navigation and mental simulation were no longer detectable within the binary classifier. When we considered individual epochs through navigation and simulation, we were now unable to classify them using standardized power.

### Standardization of low-frequency power maintains position decoding

We first sought to determine whether the standardized low-frequency power can be used to predict position during navigation to ensure that the navigation decoding results would not change. For each patient, we trained a random forest classification ensemble (see Methods) to learn position using the average standardized low frequency power (2- 12 Hz) from all hippocampal electrodes plus the identity of the start and end store as predictor variables. After standardizing power, we found that the mean accuracy for the model trained with real segment labels was still significantly greater than mean accuracy of the model trained with shuffled segment labels (Wilcoxon ranksum test p = 0.019, median real labels = 0.040, median shuffle labels = 0.025, Figure 4a). Peaks in the predictor importance for the standardized power occurred at 5.2 and 8.7 Hz).

**Figure 4:**
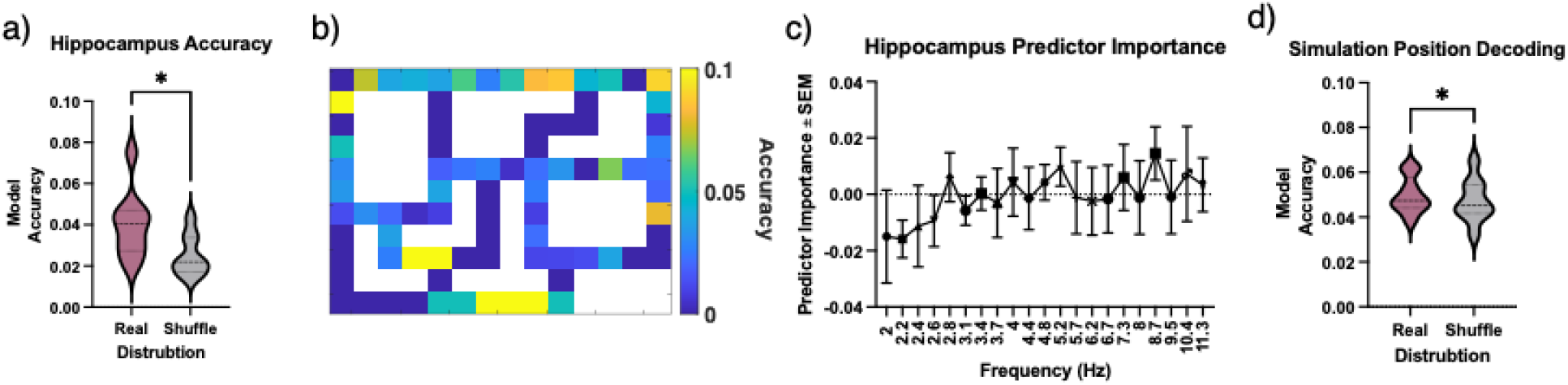
Successful position decoding navigation and mental simulation using standardized power. a. Across subjects, the mean accuracy of position decoding during navigation for the true segment labels was significantly greater than mean accuracy of the model trained with shuffled segment labels. b. Heat map showing the mean accuracy for each spatial segment. c. Average feature importance for all frequencies between 2- 12 Hz. Peaks in the predictor importance for the standardized power occurred at 5.2 and 8.7 Hz. d. Across subjects, the mean accuracy of position decoding during mental simulation using the model trained during navigation was significantly greater than mean accuracy of the model trained on the shuffled time series.

These results suggest that after standardizing the power within navigation, there is sufficient information present to decode position during navigation. When we compared the accuracy of position decoding during navigation between the model trained on raw power to the model trained on the standardized power, there was no difference in the accuracy between the two models (Wilcoxon ranksum p = 0.098, mean with raw= 0.033, mean with standardized =0.040, Supplemental Figure 3a). These findings support the idea that there are patterns within the local field potential, both in the raw and standardized power that are sufficient to decode position during navigation. These findings replicate past findings from other published work suggesting that fluctuations in the local field potential reveal information about position during navigation (Agarwal et al., 2014; Herweg et al., 2023).

### Information content of low-frequency standardized power overlaps between navigation and memory

We next tested whether we could decode position during mental simulation using the model that was trained on the standardized power during navigation, which allowed us to remove any overshadowing effects that power and frequency might have been removed. For the model trained on standardized power during navigation and applied to the standardized power during mental simulation, the mean decoding accuracy during mental simulation was significantly greater than expected by chance (Wilcoxon ranksum test p = 0.027, accuracy true timeseries = 0.050, accuracy shuffled timeseries = 0.0473, Figure 4d). In other words, the model trained during navigation could be used to decode position during mental simulation. This suggests that there is memory-related overlap in the information embedded in the low-frequency power during navigation and simulation.

To validate the finding that there is overlap in the pattern of low-frequency power during navigation and mental simulation using standardized power, we ran a correlation between the power at all time points during navigation and the power at all time points during mental simulation. We reasoned that this analysis should provide evidence of systematic increases in the correlation of power between navigation and mental simulation if position information from navigation is embedded in the local field potential recorded during mental simulation (Figure 5a, see Methods). We divided each navigation and mental simulation trial into two time-windows of equal length (early and late) and subsequently averaged the power correlation r-values between navigation and mental simulation into each of the 4 windows (early navigation to early mental simulation, early navigation to late mental simulation, late navigation to early mental simulation and late navigation to late mental stimulation). If the patterns of low-frequency power unfold in a temporally similar manner during navigation and mental simulation (albeit compressed), this implies that the power between early navigation and early mental simulation should be higher than the correlation between early navigation and late mental simulation (and the same for the second half of navigation and second half of mental simulation) (Figure 5b).

**Figure 5:**
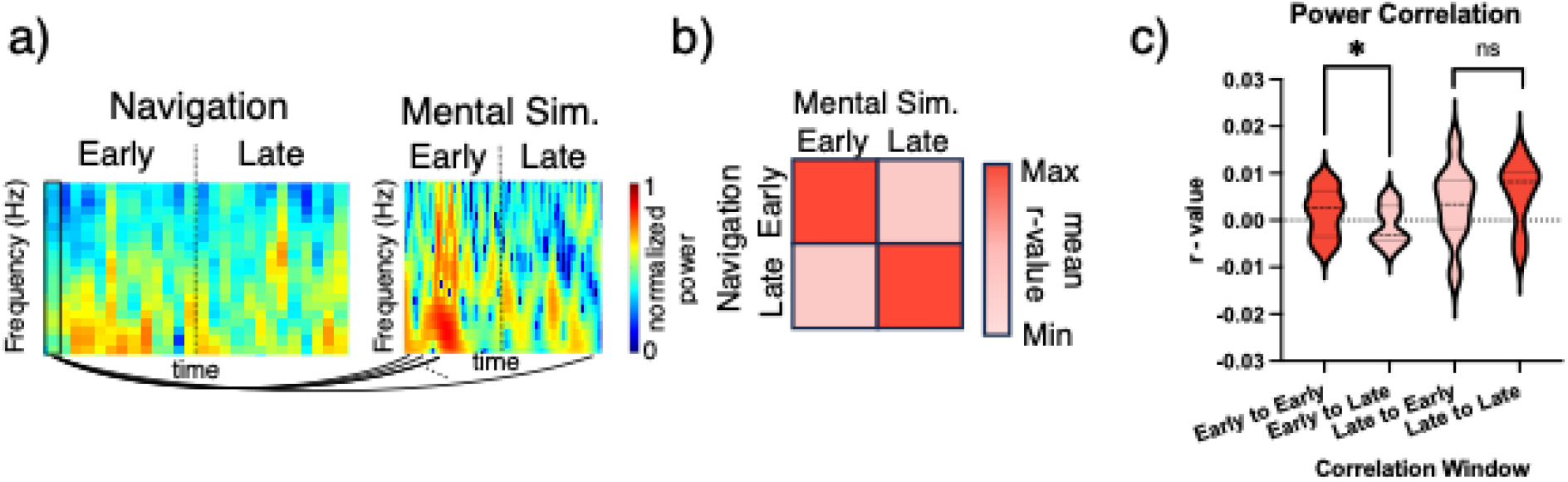
Power during early navigation and early mental simulation time points a. Power for an example navigation and mental simulation trial for a single electrode showing the correlation between navigation and mental simulation windows. Low-frequency power during navigation was averaged into 500 ms time bins, and the power at each time bin during navigation was correlated with the power at each time point during navigation. Time bins during navigation and time points during mental simulation were divided into early and late periods. b. Matrix showing the predicted correlation values for each of the four possible navigation and mental simulation correlation windows. The correlation for the two matched windows (early navigation to early mental simulation and late navigation to late mental simulation) should be greater than the correlation for the two mismatched windows (early navigation to late mental simulation and late navigation to early mental simulation). c. The correlation between early navigation and early mental simulation was significantly greater than the correlation between early navigation and late mental simulation. The correlation between late navigation and late mental simulation was not significantly greater than the correlation between late navigation and early mental simulation.

We first ran this correlation analysis using the standardized power during navigation and mental simulation to attempt to validate our previous findings showing we could decode the content of mental simulation using the hippocampal LFP recorded during navigation. A direct comparison of average correlation values across subjects revealed that the correlation in the early-early time window was significantly higher than that in the early-late window (Wilcoxon rank-sum test, p = 0.048, mean correlation r-values for early-early = 0.002, early-late = -0.001; see Figure 5c). No significant difference was observed between the late-late and late-early windows (Wilcoxon rank-sum test, p = 0.367, mean correlation r-values for late-late = 0.005, early-late = 0.003, Figure 5c). We conducted the same analysis using raw power and did not find any significant differences (early-early vs. early-late, Wilcoxon rank-sum test, p = 0.082, mean correlation r-values for early-early = 0.592, early-late = 0.601, see Supplemental Figure 4a; late-late and late-early windows: Wilcoxon rank-sum test, p = 0.455, mean correlation r-values for late-late = 0.595, early-late = 0.597; Supplemental Figure 4a). The lack of a significant difference in the strength of the correlation between any matched windows supports the conclusion that the differences in the raw power between navigation and mental simulation obscures subtle patterns in the local field potential. The finding that the correlation is only significant for matched early time periods converges with the time-resolved decoding analysis showing the highest accuracy at the start of mental simulation (Supplemental Figure 3c), suggesting that the overlap in patterns of low-frequency power between navigation and mental simulation occur during early time periods of mental simulation. Overall, these findings align with the hypothesis that there are overlapping low-frequency power dynamics during navigation and mental simulation that can be detected after accounting for differences in the underlying power spectra. Yet, both the classifier and correlation analyses suggest that these are relatively weak signals embedded within the hippocampal LFP and the overlap is strongest between early portions of navigation and mental simulation.

### Position decoding during mental navigation is specific to the hippocampus

We repeated the same analysis from above on electrodes that were located in either the frontal cortices or parietal cortices (see Methods). During navigation, we found that the model accuracy for decoding position was significantly greater than expected by chance for both the parietal cortices (Wilcoxon ranksum test p = 0.008, median true labels = 0.040, median shuffle labels = 0.022, Figure 6a) and frontal cortices (Wilcoxon ranksum test p = 0.020, median true labels = 0.039, median shuffle labels = 0.021, Figure 6b). This result shows that the ability to use low-frequency power to decode position during navigation is not specific to the hippocampus.

**Figure 6:**
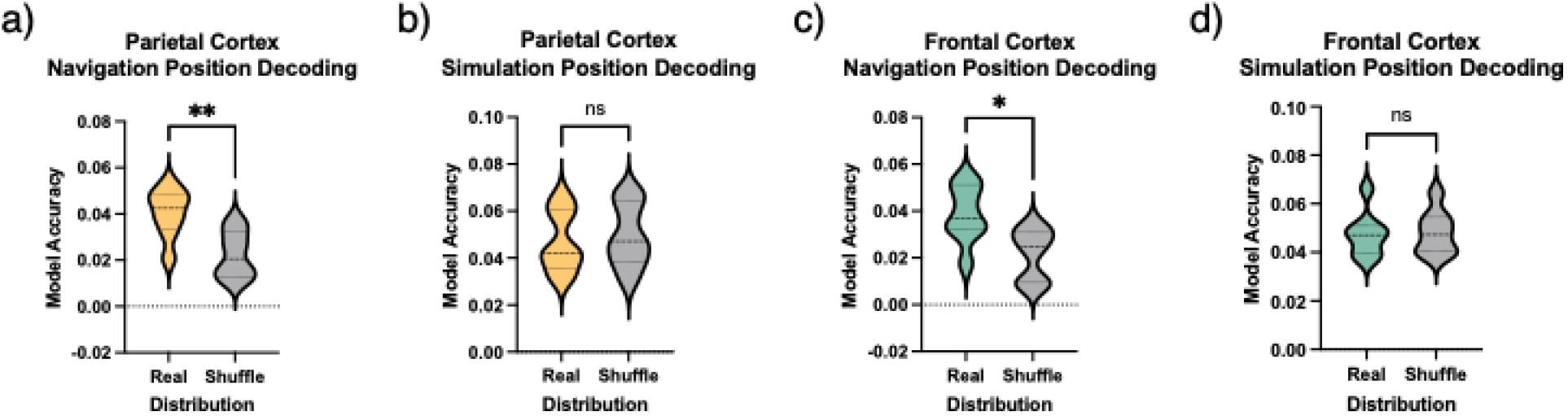
Only navigation position is successful in the frontal and parietal cortices a. When using electrodes localized to the parietal cortex, the mean accuracy of position decoding during navigation for the true segment labels was significantly greater than mean accuracy of the model trained with shuffled segment labels across subjects. b. When using electrodes localized to the frontal cortex, the mean accuracy of position decoding during navigation for the true segment labels was significantly greater than mean accuracy of the model trained with shuffled segment labels across subjects. c. When using electrodes in the parietal cortex, the mean accuracy of position decoding during mental simulation using the model trained during navigation was not significantly greater than mean accuracy of the model trained on the shuffled time series. d. When using electrodes in the frontal cortex, the mean accuracy of position decoding during mental simulation using the model trained during navigation was not significantly greater than mean accuracy of the model trained on the shuffled time series.

When we applied the model to mental simulation, we found that there was no significant difference in accuracy for the true and the shuffled timeseries (Wilcoxon ranksum test p = 0.125, median true timeseries = 0.046, median shuffle timeseries = 0.049, Figure 6c) nor the frontal cortices (Wilcoxon ranksum test p = 0.273, median true timeseries = 0.047, median shuffle timeseries = 0.048, Figure 6d) . This result implies that overlap in the content embedded in low frequency power during navigation and mental simulation was specific to the hippocampus.

## Discussion

The key finding of this study is that the position of an imagined route during mental simulation can be decoded using a classifier trained on standardized local field potential (LFP) recordings collected during navigation. This effect was specific to the hippocampus and absent in recordings from the parietal and frontal cortex. These results suggest that memory-related information embedded in the LFP during navigation is sufficiently preserved during mental simulation, enabling above-chance decoding of the participant’s simulated position. To validate this finding, we examined the correlation between LFP patterns during navigation and mental simulation. We observed significantly higher correlations during matched early stages of both tasks compared to mismatched early and late stages. Additionally, a time-resolved decoding analysis revealed that the content of mental simulation is most decodable at the beginning of simulation and nearly absent in later stages. Together, these results underscore the hippocampus’ critical role in memory-related processing, particularly its dual function in episodic memory and the replay of content during mental simulation.

A notable caveat of our analysis is that position decoding during mental simulation was only successful with standardized LFP data, which accounts for baseline power spectra differences. Using the raw LFP data, we could not decode simulated positions, although the position of the participant remained decodable. This highlights the importance of within-condition power normalization, as differences in absolute power during navigation and simulation otherwise obscure overlapping patterns across electrodes. The observed differences in low-frequency power between navigation and simulation align with previous findings indicating stronger theta oscillations during mental simulation than during spatial navigation (Seger et al., 2023). These findings suggest that memory replay is a more prominent driver of hippocampal theta than navigation. By normalizing background power differences, we were able to uncover weak but significant signals common to both navigation and simulation. This supports the idea that internally generated dynamics during mental simulation are potent drivers of theta activity. Furthermore, despite these differences, overlapping memory-related signals in the LFP can still be detected across both conditions.

Considerable debate remains about whether the hippocampus can best be considered a brain structure devoted to spatial navigation or episodic memory (Eichenbaum, 2017a). We believe that the findings here (and previous results (Seger et al., 2023) help to shed light on this important issue. In our previous paper, we found that theta power and frequency were stronger during episodic memory and mental simulation of the same route than during navigation. This would support the idea that the hippocampus plays an important role in internally generated dynamics related to episodic memory (Miller et al., 2013; Wang et al., 2015; Vass et al., 2016; Wanjia et al., 2021; Zou et al., 2023) rather than navigation. Navigation, however, involves some memory-related components (Ekstrom and Hill, 2023), and during navigation, participants encoded the route they would then simulate. Therefore, by demonstrating that there is sufficient information in the standardized local field potential to decode position from mental simulation based on navigation, our result support the idea that the role of the hippocampus during navigation is likely related to memory more so than navigation (Eichenbaum, 2017a, 2017b).

It has been found that place cells during navigation are reactivated during episodic memory retrieval; however, it is unclear if this reactivation is accompanied by similar patterns in the low-frequency oscillatory patterns. Previous papers have investigated periods of replay using brain recordings during encoding and retrieval, including list learning and learning of visual associations (Michelmann et al., 2016, 2019; Wimmer et al., 2020; Crivelli-Decker et al., 2023). These papers employed either scalp EEG, MEG, or fMRI and therefore were not able to specifically localize time-resolved signals to the hippocampus. Another potential limitation with some of these previous studies is that they did not have a behavioral measure to ensure that participants were replaying specific content from encoding. Here, we overcame this issue by using reaction time as a proxy for route distance and including catch trials that ensured participants were simulating the route. Finally, these papers did not involve comparison with navigation through physical space, and it is possible that some of the signal in these studies was related broadly to episodic memory rather than specific contents of previously encoded memories. Our results provide an important advance by showing that the engagement of the hippocampus during navigation and simulation likely relates most strongly to encoding and retrieval of episodic content.

Another finding from our study is that we could successfully decode the position of the participants when navigating in a complex environment using both the raw and standardized LFP. Previous studies have also shown that it is possible to decode position from the local field potential in both rats (Cao et al., 2021) and humans (Watrous et al., 2011; Graves et al., 2023; Herweg et al., 2023). Our findings here replicate this basic result and extend it by showing that this is also true in more complex spatial environments and that such a signal is also present in parietal and frontal cortices. These findings suggest that position is coded in a distributed manner in the human brain during navigation and not specific to the hippocampus, consistent with some previous reports (Sargolini et al., 2006; Ji and Wilson, 2007; Miller et al., 2013; Jankowski and O’Mara, 2015; Mao et al., 2017; Sauer et al., 2022; Du et al., 2023). We also showed that this position-related signal was largely restricted to the lower frequencies of the local field potential, primarily centered within the human theta band (4 – 8 Hz). While theta oscillations appear to be ubiquitous across many different behaviors, our findings, coupled with previous reports, support the idea that there are subtle fluctuations in the low-frequency bands that are specific to position. It is not clear if these fluctuations in low-frequency power relate to potential differences in position-specific attention or some heretofore undiscovered cognitive phenomenon that might be coded in the local field potential and relate to spatial memory for position. Nonetheless, these fluctuations in the local field potential, once we had removed overshadowing effects of power and frequency, were also decodable when the same route was simulated following navigation. These findings overall support the idea that human theta oscillations support memory-related content. While this memory-related content is present during both navigation and mental simulation of the same route, our results suggest that memory-related content is likely the driver of theta oscillations during navigation rather than sensorimotor-related components.

## Methods

### Behavioral Methods

Patients performed a spatial navigation task on a laptop computer using an Xbox controller. The task was designed in Unity 3D using the “Landmarks” framework from GitHub (Starrett et al., 2021)(). In this task, participants navigated routes between stores located within a sparse environment and mentally simulated the routes immediately upon route completion. The task was designed as follows. Before entering the virtual environment, the participant completed the Storefront Familiarization portion of the task during which the six storefronts were displayed successively on a blank screen for 5000 milliseconds(ms) to familiarize the participants with the target stores. Between the appearance of each storefront a white crosshair was displayed on a black background for 5000ms. After viewing each storefront 2x, the participant began the Navigation Only portion of the task.

The Navigation Only portion of the task began when the participant appeared outside one of the stores in the virtual environment in a first-person point of view, and a semi-transparent on-screen prompt instructed the participant to locate one of the five remaining target stores. The prompt remained overlaid on the entire screen for 5000ms, during which time the participant was able to begin using the XBox joystick to navigate to the target store. After 5000ms, the on-screen prompt was reduced in size and moved to the top right corner of the screen. Once the participant reached the target store, another semi-transparent on-screen prompt was overlaid on the screen for 5000 ms indicating that the target store was identified. During Navigation Only, the prompt indicating the identification of the target store was followed by a white crosshair that was displayed on a black background for 5000ms (e.g. the environment was not visible). Following the crosshair, the participants’ previous view of the environment was restored, and a new prompt instructed the participant to locate the next target store.

After the participant visited each store 2x, the participant began the Navigation with Mental Simulation portion of the task during which the participant was instructed to “mentally navigate” the route upon arrival at the target store (See Instructions for Mental Simulation). The task structure of Navigation with Mental Simulation was similar to Navigation Only; however, the on-screen prompt that indicated the successful identification of the target store was instead followed by a self-paced on-screen prompt which instructed the participant to press [A] on the controller to begin mental simulation of the route and [A] again when they were done. When the participant pressed [A] to begin mental simulation, the instructions on the screen were replaced by a white crosshair, which the participants were instructed to look at while actively mentally navigating. When the participant pressed [A] to indicate the end of mental simulation, the previous view of the environment was restored and a new on-screen prompt overlaying the environment instructed the participant to locate the next store. This procedure was repeated 44x.

For each of the 44 trials, the participant was assigned to one of two speeds—5 vm/s (slow speed) or 8 vm/s (fast speed). On trials in which the participant navigated without mental simulation, the speed of movement was equal to 6 vm/s (medium speed). 10% of the 44 navigation + mental simulation trials consisted of catch trials. The speed manipulation was not considered in this manuscript. On catch trials, the participant was instructed to use the controller to simulate the imagined route during mental simulation. The catch trials were to ensure that participants were actually remembering the route when mentally simulating.

Instructions for Mental Simulation and Practice Task After consenting to the task and prior to starting, each participant completed a practice exercise to become familiar with the task design. The practice exercise consisted of asking the participant to imagine standing at home in their bedroom and then imagine walking from this location in the bedroom to the kitchen. All participants were successfully able to complete this exercise before beginning the task.

At the beginning of the task, the participant completed a practice version of the entire task. The practice version of the task was identical to the full version; however, the number of trials was condensed (2 navigation trials, 2 navigation + mental simulation trials) and the virtual environment consisted of 2 stores located on either end of a straight road. There were no catch trials during mental simulation in the practice version of the experiment. The practice version was designed to familiarize participants with the task structure and instructions for mental simulation prior to the experiment.

### Patient Data

10 patients with medically intractable epilepsy who underwent stereo- electroencephalography surgery for clinical purposes were recruited to participate in this study. In total, there were 6 males and 4 females between 21 - 63 years of age (median 41 years old). Data were collected from the University of Texas Southwestern Medical Center epilepsy program. Electrode placement was dictated solely by the clinical need for seizure localization. Each subject was implanted with up to 17 depth electrodes containing 8-16 cylindrical platinum– iridium recording contacts spaced 2-6-mm apart. Following implantation, electrode localization was achieved by co-registration of the post-operative computed tomography scans with the pre- operative magnetic resonance images using the FSL’s/FLIRT software(41). The co-registered images were evaluated by an expert member of the neuroradiology team to determine the final electrode locations. Each subject had at least three recording contacts localized to the hippocampus. The median number of hippocampal contacts per subject was 6. The median number contacts in the anterior hippocampus per subject was 3.5, and the median number in the posterior hippocampus per subject was 2.

The number of electrodes in the parietal cortex varied between 2 and 10, and the median number of electrodes was 8 (one subject only had 1 electrode). The number of electrodes in the frontal cortex varied between 2 and 10, and the median number of electrodes was 10.

### iEEG Data Analysis

Stereo-EEG data were recorded using a Nihon Kohden EEG-1200 clinical system. Signals were sampled at 1000 Hz and referenced to a common intracranial contact. Raw signals were subsequently re-referenced to a common average montage, with each contact referenced to the average across all contacts and then down sampled to 250Hz. All analyses were conducted using MATLAB with both built-in and custom-made scripts. We employed an automated artifact rejection algorithm to exclude interictal activity by computing kurtosis over a sliding window of 1000 ms duration and a kurtosis threshold of 5.

### Environment Segmentation

The walkable area of the spatial environment was divided into 82 equally spaced bins with a size of 10x10 vm, as shown in Figure 1c.

### Power Spectral Density Estimation

Each participant’s EEG data was filtered with a 6th-order highpass Butterworth filter at with a cutoff frequency of 1Hz. Power was estimated for log-spaced frequencies between 2-12Hz using a continuous wavelet Morlet transform (width = 5) for every time point during. each navigation and mental simulation trial for every electrode. A 2-second buffer was added to avoid edge artifacts and removed after extracting power.

In additional to extracting the raw power for all trials, a standardized version of the power was also created for all navigation and simulation trials with the goal of reducing the differences in the power values between navigation and mental simulation while still maintaining relevant information. To do this, we computed the mean and standard deviation of power for across all navigation and mental simulation time points (separately for each condition). For each time point during navigation and mental simulation, we subtracted the respective means and divided by the standard deviation.

To compute the average power for the segments traversed during a single trial, the position at each time point was assigned to a spatial bin and the power was average across all time points for each frequency. This process was repeated for all trials to yield a matrix of the average power at each frequency for all segments traversed throughout the trial. This process was repeated for all electrodes.

In addition to segmenting position into spatial bins, time points during navigation were also broken down into 5 conditions (regardless of environment position): start (first 1 second of navigation), end (last 1 second of navigation) straight (participant is moving along a straight road segment), immobile (time periods of no movement), forced turns (participant turns due to bend in road), optional turn (participant chooses to make a turn at an intersection), and preceding optional turns (1 second period prior to an optional turn). To compute the average power for each condition for each navigation trial, the position at each time point was assigned to one of the 5 conditions. For each trial, the power was averaged across all time points at each frequency for each of the 5 conditions. This process was repeated for all trials and all electrodes. To compute the average power during the simulation condition, for each trial we randomly selected a 1 second interval (excluding the first and last 1 second). For each simulation trial, the power was averaged across all time points at each frequency. This process was repeated for all trials and all electrodes.

### Theta Frequency Range and Bands

We defined the theta band as frequencies from 2 - 12 Hz because this was roughly the range over which the PSD differed for navigation and mental simulation (Seger et al., 2023)

### Random Forest Analysis

#### Position

To generate training data for each subject, the average power for all segments traversed during navigation was combined across all electrodes. A single sample for a given segment consisted of the average power at each frequency for all electrodes for a given subject. After combining power across electrodes, any segments that had less than 15 samples were removed from the training data. The number of samples per segment varied between 15 – 30, depending on the subject and segment. To ensure an equal number of training samples for each segment, we randomly selected 15 samples for inclusion in training. In order to sample all of the available data, we repeated the random sample selection process to generate 10 training data sets with different a combination of samples in each. The number of segments that were included in the training dataset (i.e. number of segments with 15 samples) varied between 16 to 41 between subject, and the average number of segments was 29. Given the average number of distinct segments was 29, the probability of randomly correctly identifying a segment was 0.03.

Using MATLABS “fitcensemble” function, a classification ensemble with 100 learners was trained to learn segment labels using the power at each frequency for all electrodes and the label of the start and end store as predictor variables. There are 21 log-spaced frequencies sampled between 2-12 Hz and the number of electrodes localized to the hippocampus varied between 3 – 13 for each subject, meaning the number of predictors for a given subject varied between 23 – 301. For all models, feature selection was optimized based on interaction-curvature measures. To compute the model accuracy and feature importance, we used out-of-bag (OOB) predictions to measure of model accuracy and feature importance without having to split the training data into train and test sets. For each subject, a separate model was trained for each of the 10 datasets, and the accuracy and predictor importance were the averaged across all models to generate a single estimate of accuracy and predictor importance for each subject.

For each of the 10 training datasets for each subject, a shuffled version of training data was also created by shuffling the segment labels. In order to estimate chance accuracy, a bagged classification ensemble was separately trained to learn segment labels using the shuffle dataset. The accuracy and predictor importance for each of the shuffled datasets were averaged across all models to generate a single estimate of chance accuracy and predictor importance for each subject.

To evaluate the model during mental simulation, for each trial we first generated a set of “expected” position locations based on the position during navigation for that trial. Specifically, we down sampled the position information during navigation so that the length of the position vector matched the length of mental simulation. The “expected” position (i.e. segment label) during at each simulation time point was then determined by the segment label from the down- sampled position vector. We then used the model trained on the true segment labels to determine the predicted position at each time point during mental simulation. Model accuracy during mental simulation was defined as the proportion of time points where the expected position and predicted position are the same. To determine the chance accuracy, the same process was used except that the predicted position at each time point was determined using model that was trained on the shuffled dataset.

To determine the accuracy of decoding at each time point during mental simulation, for each trial we took the binary vector representing the accuracy at each time point and resampled this vector to a normalized length of 1000 samples for all trials. We then computed the average accuracy at each of the 1000 normalized time points to measure the average accuracy at each time point during mental simulation.

#### Condition

Generation of the training data and model training as described below was repeated separately for each of the 6 navigation conditions of interest (start, end, straight, immobile, forced turns, optional turns).

To generate training data, the average power for one of the 6 the navigation conditions for was combined with the average power for the simulation condition.

The number of samples per condition (1 navigation and 1 simulation) varied between 30 – 40, depending on the subject and condition. To ensure equal number of samples for both conditions, we randomly selected a subset of samples from the condition with more samples to equal the number of samples in the condition with less overall samples. In order to sample all of the available data, we repeated the random sample selection process to generate 10 training data sets with different a combination of samples in each.

Using MATLABS “fitcensemble” function, a classification ensemble with 100 learners was trained to learn condition labels using the power at each frequency for all electrodes and the label of the start and end store as predictor variables. There are 21 log-spaced frequencies sampled between 2-12 Hz and the number of electrodes localized to the hippocampus varied between 3 – 13 for each subject, meaning the number of predictors for a given subject varied between 23 – 301. For all models, feature selection was optimized based on interaction-curvature measures. To compute the model accuracy and feature importance, we used out-of-bag (OOB) predictions to measure of model accuracy and feature importance without having to split the training data into train and test sets. For each subject, a separate model was training for each of the 10 datasets, and the accuracy and predictor importance were the averaged across all models to generate a single estimate of accuracy and predictor importance for each subject.

For each of the 10 training datasets for each subject, a shuffled version of training data was also created by shuffling the condition labels. In order to estimate chance accuracy, a bagged classification ensemble was separately trained to learn segment labels using the shuffle dataset.

The accuracy and predictor importance for each of the shuffled datasets were averaged across all models to generate a single estimate of chance accuracy and predictor importance for each subject.

### Power Correlations

For each subject, for each subject we divided each navigation trial into 500 ms time bins and computed the average power at each frequency for each electrode. For each of the 500 ms bins, we concatenated the average power for each frequency across all electrodes for each subject to generate a 1xN matrix (number of electrodes * number of frequencies) for each 500 ms bin. We then took the power at each mental simulation time point and combined across electrodes to generate a 1xN matrix (number of electrodes * number of frequencies) at each mental simulation time. We ran a Spearman correlation between each navigation time bin and all mental simulation time points for all trial pairs to estimate the correlation between each time point during navigation and each time point during mental simulation. We divided each navigation and mental simulation trial into early and late periods (first half of time points in the trial are considered ‘early’ and second half of time points are considered ‘late’). For each trial, we then averaged the correlation values into four groups based on—early navigation and early mental simulation correlations (early-early), early navigation and late mental simulation correlations (early-late), late navigation and early mental simulation correlations (late-early), and late navigation and late mental simulation correlations (late-late).

### Statistics

To compare model accuracy for the decoding of position and condition, we ran a one-tailed paired Wilcoxon ranksum test between the subject level average accuracy for model trained on the dataset with true labels and the accuracy of the model trained in the dataset with shuffled data.

For the correlation analysis, we performed a 2-way ANOVA to compare the effect of navigation and mental simulation time windows on the strength of the correlation. To determine whether the correlation for the ‘early-early’ window was greater than’ early-late’ window we ran a one-tailed paired Wilcoxon ranksum test between the average correlation for ‘early-early’ windows and ‘late-late’ windows across subjects. The same process was repeated for the same for ‘late-late’ and ‘late-early’ windows.

