## Supplemental Figures for "Differences and similarities between human hippocampal low-frequency oscillations during navigation and mental simulation"

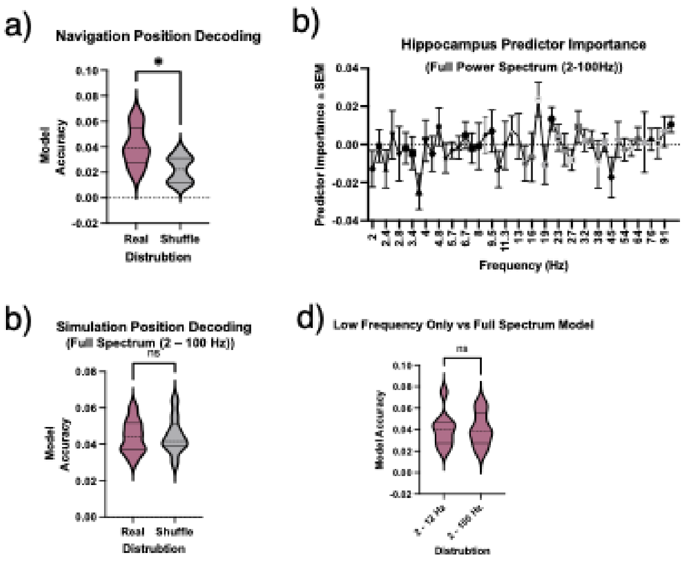


Supplemental Figure 1:

a. When all frequencies between 2 – 100 Hz were included, the mean accuracy of position decoding during navigation for the true segment labels was significantly greater than mean accuracy of the model trained with shuffled segment labels (Wilcoxon ranksum test p = 0.020, median real labels = 0.040, median shuffle labels = 0.022).
b. Average feature importance for all frequencies between 2- 100 Hz.
c. When all frequencies between 2 – 100 Hz were included, the mean accuracy of position decoding during mental simulation using the model trained during navigation was not significantly greater than mean accuracy of the model trained on the shuffled time series (Wilcoxon ranksum test p = 0.362, accuracy  true timeseries = 0.044, accuracy shuffled timeseries = 0.045).
d. There was no significant difference between decoding accuracy for the model trained using power for frequencies 2 – 12 Hz compared to the model trained on using 2 – 100Hz power. (Wilcoxon ranksum test p = 0.999, accuracy 2- 12 Hz = 0.040, accuracy 2 – 100Hz = 0.040).

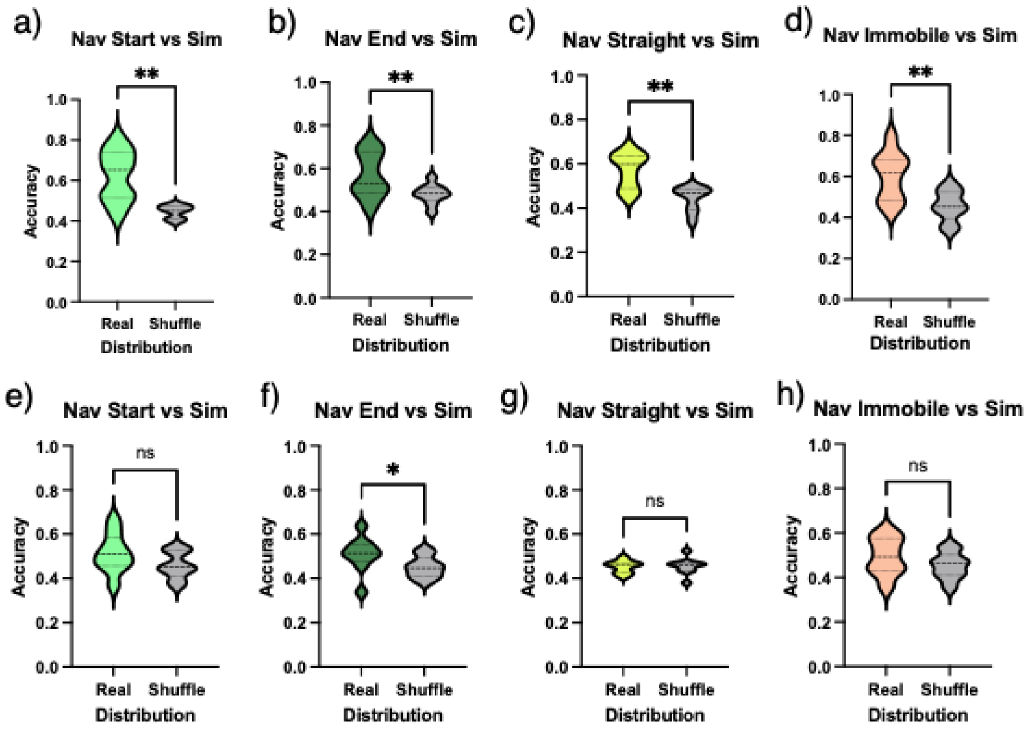


Supplemental Figure 2:

a. When using the raw low-frequency power, the mean accuracy for decoding navigation (start only, see Methods) versus mental simulation condition for the model trained with true condition labels was significantly greater than the model trained with shuffled condition labels across all subjects (Wilcoxon ranksum test p = 0.008, mean true accuracy = 0.623, mean shuffle accuracy = 0.445).
b. When using the raw low-frequency power, the mean accuracy for decoding navigation (end only, see Methods) versus mental simulation condition for the model trained with true condition labels was significantly greater than the model trained with shuffled condition labels across all subjects (Wilcoxon ranksum test p = 0.008, mean true accuracy = 0.569, mean shuffle accuracy = 0.480).
c. When using the raw low-frequency power, the mean accuracy for decoding navigation (straight only, see Methods) versus mental simulation condition for the model trained with true condition labels was significantly greater than the model trained with shuffled condition labels across all subjects (Wilcoxon ranksum test p = 0.008, mean true accuracy = 0.571, mean shuffle accuracy = 0.444).
d. When using the raw low-frequency power, the mean accuracy for decoding navigation (immobile only, see Methods) versus mental simulation condition for the model trained with true condition labels was significantly greater than the model trained with shuffled condition labels across all subjects (Wilcoxon ranksum test p = 0.004, mean true accuracy = 0.601, mean shuffle accuracy = 0.456).
e. When using the standardized low-frequency power, the mean accuracy for decoding navigation (start only, see Methods) versus mental simulation condition for the model trained with true condition labels was not significantly greater than the model trained with shuffled condition labels across all subjects (Wilcoxon ranksum test p = 0.054, mean true accuracy = 0.513, mean shuffle accuracy = 0.459).

f. When using the standardized low-frequency power, the mean accuracy for decoding navigation (end only, see Methods) versus mental simulation condition for the model trained with true condition labels was significantly greater than the model trained with shuffled condition labels across all subjects (Wilcoxon ranksum test $p = 0.027$, mean true accuracy = 0.509, mean shuffle accuracy = 0.451).
g. When using the standardized low-frequency power, the mean accuracy for decoding navigation (straight only, see Methods) versus mental simulation condition for the model trained with true condition labels was not significantly greater than the model trained with shuffled condition labels across all subjects (Wilcoxon ranksum test $p > 0.999$, mean true accuracy = 0.453, mean shuffle accuracy = 0.455).
h. When using the standardized low-frequency power, the mean accuracy for decoding navigation (immobile only, see Methods) versus mental simulation condition for the model trained with true condition labels was not significantly greater than the model trained with shuffled condition labels across all subjects (Wilcoxon ranksum test $p = 0.250$, mean true accuracy = 0.486, mean shuffle accuracy = 0.494).

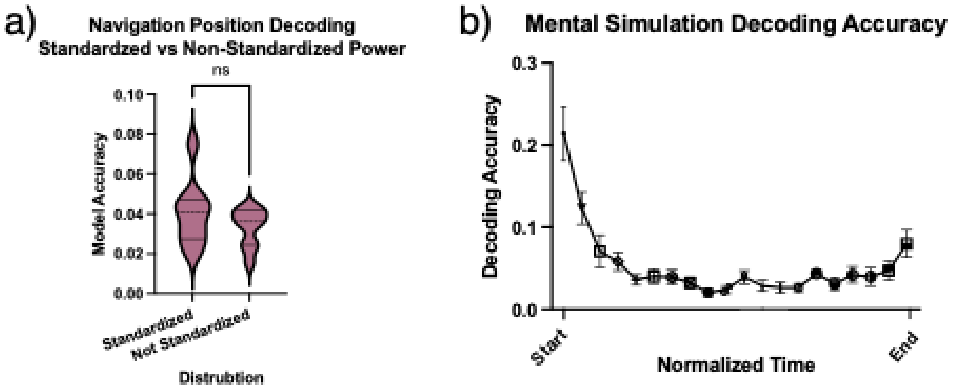


Supplemental Figure 3: Decoding position during mental simulation using the model trained during navigation on the standardized 2 – 12 Hz power.

a. There was no significant difference between decoding accuracy during navigation for the model trained using the raw 2 – 12 Hz power compared to the model trained using thee standardized 2- 12 Hz (Wilcoxon ranksum p = 0.098, mean with raw= 0.033 , mean with standardized = 0.040).
b. Mean decoding accuracy during mental simulation as a function of normalized time (see Methods).

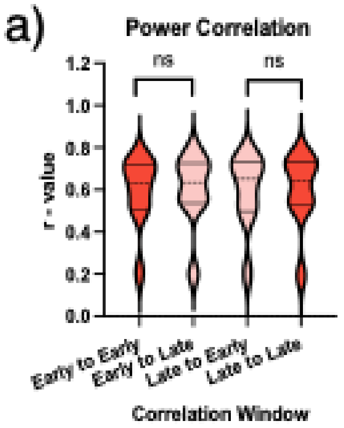


Supplemental Figure 4:

a. When using the raw power, the correlation between early navigation and early mental simulation was not significantly greater than the correlation between early navigation and late mental simulation (Wilcoxon rank-sum test, $p = 0.082$, mean correlation r-values for early-early = 0.592, early-late = 0.601). The correlation between late navigation and late mental simulation was not significantly greater than the correlation between late navigation and early mental simulation (Wilcoxon rank-sum test, $p = 0.455$, mean correlation r-values for late-late = 0.595, early-late = 0.597).

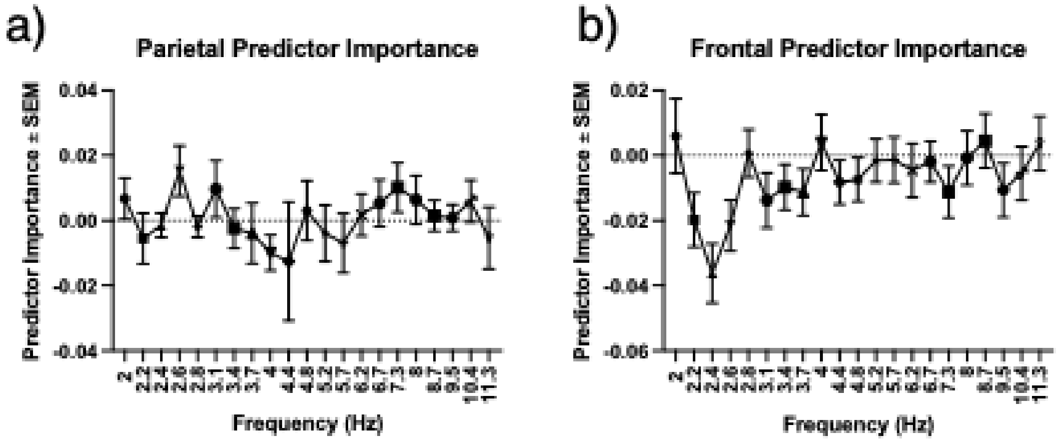


Supplemental Figure 5:

a. Average feature importance for all frequencies between 2- 12 Hz for electrodes in the parietal cortex.
b. Average feature importance for all frequencies between 2- 12 Hz for electrodes in the frontal cortex.
